# Regulation of Numb during planar cell polarity establishment in the Drosophila eye

**DOI:** 10.1101/686840

**Authors:** Pedro M Domingos, Andreas Jenny, David del Alamo, Marek Mlodzik, Hermann Steller, Bertrand Mollereau

**Affiliations:** Instituto de Tecnologia Química e Biológica, Universidade Nova de Lisboa, Av. da República, Oeiras 2780-157, Portugal; Strang Laboratory of Apoptosis and Cancer Research, The Rockefeller University, Box 252, 1230 York Avenue. New York, NY 10065, USA; Albert Einstein College of Medicine, 1300 Morris Park Avenue, Chanin Building, Room 503, Bronx, NY10461, USA; Department of Cell, Developmental, and Regenerative Biology, Icahn School of Medicine at Mount Sinai, One Gustave L. Levy Place, New York, NY 10029, USA; European Molecular Biology Organization, Meyerhofstrasse 1, 69117 Heidelberg, Germany; Université de Lyon, ENSL, UCBL, CNRS, LBMC, 46 Allée d’Italie, 69007, Lyon, France

**Keywords:** *Drosophila* eye development, Planar Cell Polarity, numb, fat

## Abstract

The establishment of planar cell polarity (PCP) in the *Drosophila* eye requires correct specification of the R3/R4 pair of photoreceptor cells, determined by a Frizzled mediated signaling event that specifies R3 and induces Delta to activate Notch signaling in the neighboring cell, specifying it as R4. Here, we investigated the role of the Notch signaling negative regulator Numb in the specification of R3/R4 fates and PCP establishment in the *Drosophila* eye. We observed that Numb is transiently upregulated in R3 at the time of R3/R4 specification. This regulation of Numb levels in developing photoreceptors occurs at the post-transcriptional level and is dependent on Dishevelled, an effector of Frizzled signaling, and Lethal Giant Larva. We detected PCP defects in cells homozygous for *numb^15^*, but these defects were due to a loss of function mutation in *fat* (*fat^Q805*^*) being present in the *numb^15^* chromosome. However, mosaic overexpression of Numb in R4 precursors (only) caused PCP defects and *numb* loss-of-function had a modifying effect on the defects found in a hypomorphic *dishevelled* mutation. Our results suggest that Numb levels are upregulated to reinforce the bias of Notch signaling activation in the R3/R4 pair, two post-mitotic cells that are not specified by asymmetric cell division.

## INTRODUCTION

Planar cell polarity (PCP) occurs when epithelial cells are polarized along the plane of the epithelium (perpendicular to the apical-basal axis of the cell). Although PCP is evident in all organisms and many different biological systems – these can include coordinated orientation of bristles in invertebrates, sensory cell orientation in the mammalian inner ear, scales in fish, or feathers in birds – and has also been linked to diseases ranging from ciliopathies to cancer, it is most studied and by far the best understood in *Drosophila* (Singh and Mlodzik, 2012) (Matis and Axelrod, 2013) (Butler and Wallingford, 2017) (Humphries and Mlodzik, 2018).

In the *Drosophila* eye, PCP is established upon specification of the R3 and R4 photoreceptors, and the subsequent rotation movement performed by the developing ommatidial clusters (Mlodzik, 1999) (Strutt and Strutt, 1999). In the adult eye, each ommatidium contains six outer PRs (R1-R6), which are positioned in a trapezoidal arrangement, with the two inner R7 and R8 located in the center of this trapezoid. The ommatidial trapezoidal arrangement comes in two chiral shapes, generated through the asymmetric positioning of R3 and R4, that form a symmetric image on either side of the dorso-ventral (D/V) midline, the equator (Fig. 1).

**Figure 1 -.**
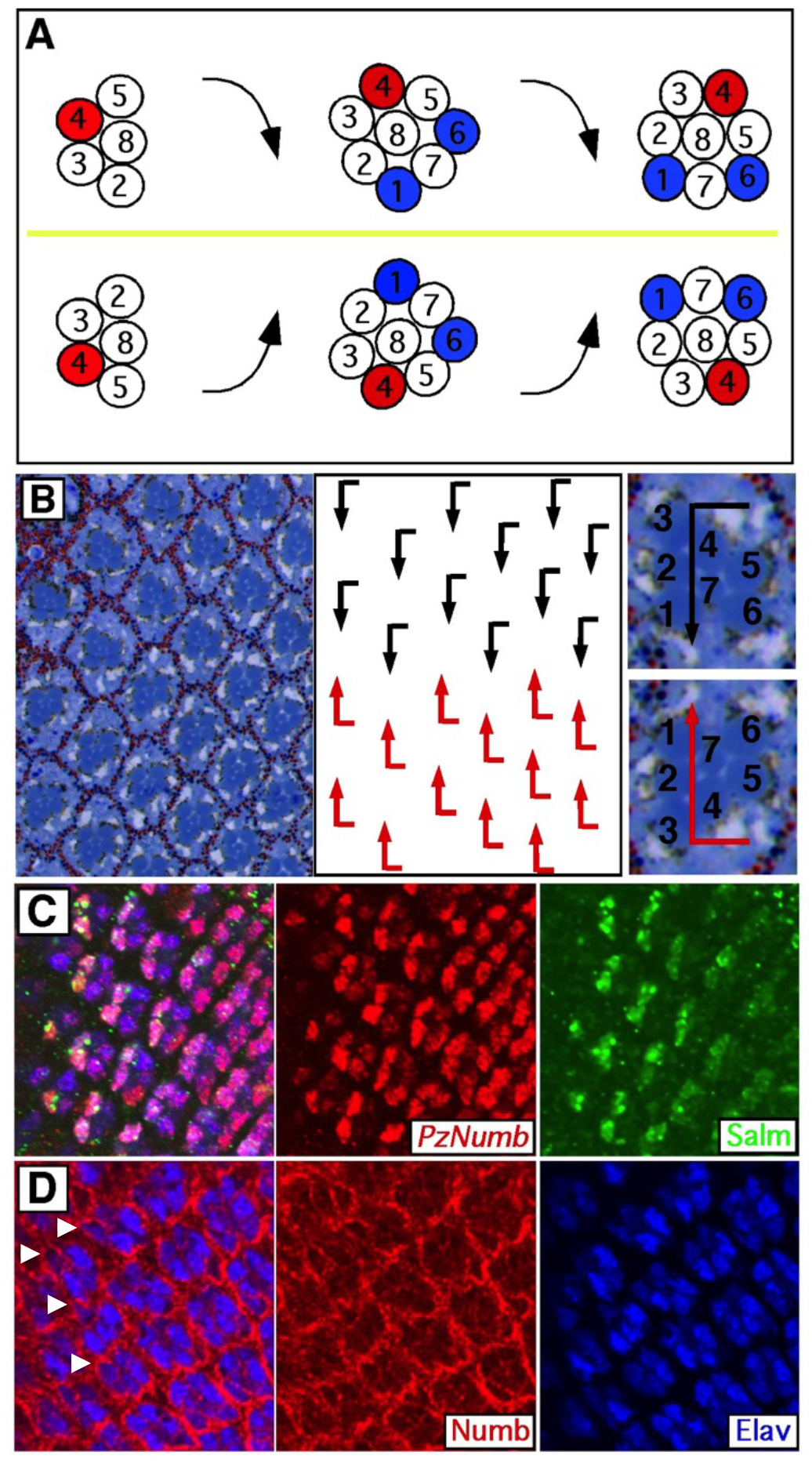
Numb is expressed in R3 and R4 when planar cell polarity is established. (A) Schematic drawing illustrating ommatidial rotation in the 3rd larval instar imaginal disc. Closer to the morphogenetic furrow (to the left) the R3/R4 pair is perpendicular to the dorsal/ventral midline, the equator (yellow line). Of the R3/R4 pair, the cell farther from the equator activates Notch signaling, starts to express *mδ0.5-lacZ* (red) and acquires the R4 fate, concomitants with an initial, fast, 45° rotation of the ommatidia, in a clockwise direction in the dorsal half and counter-clockwise direction in the ventral half of the eye imaginal disc. In the more posterior part of the eye disc (to the right), the 90° rotation of the ommatidia is complete, and the R3/R4 pair is parallel to the equator. The expression of BarH1 in R1/R6 (blue) allows the visualization of the rotation of the ommatidia. (B) Tangential section of an adult eye (left), corresponding schematic drawing (center) and single dorsal and ventral ommatidia (right). Due to R3/R4 specification and ommatidial rotation in the larval eye disc, in the adult eye ommatidia are arranged as two opposite chiral forms separated by the equator. Ommatidia in the dorsal half are represented with black arrows and in the ventral half with red arrows. Magnification of one dorsal (top) and one ventral (bottom) ommatidium with respective arrows. The numbers indicate the identities of each photoreceptor. (C) Eye imaginal disc stained for Salm (green) and lacZ (*P(PZ)numb^03235^*-red). The expression of Numb in R3/R4 is concomitant with Salm. D) Numb expression (red), visualized with anti-Numb antibody, is transiently stronger in R3 (arrowheads) than R4, when R3/R4 are specified. Elav (blue) is a marker of the photoreceptors. In this and subsequent figures, anterior is to the left and dorsal is up in all panels. Scale bars represent 10 μm.

In 3rd larval instar, the developing ommatidial clusters rotate 90° in opposite directions in the dorsal and ventral halves of the eye (clockwise in the dorsal and anticlockwise in the ventral; Fig. 1). The direction of rotation and the chirality adopted by the ommatidia are a direct consequence of the specification of the R3/R4 pair. The cell of the R3/R4 precursor pair, which is closer to the equator, adopts the R3 fate, while the other cell of the pair takes on the R4 fate. R3 specification is mediated by the activation of the Frizzled/Planar Cell Polarity (Fz/PCP) pathway in the R3 precursor and directed asymmetric activation of Notch signaling in the neighboring R4 precursor, leading to its specification as R4. Gain and loss of function experiments with several members of the Fz/PCP core group or the Notch signaling pathway lead to random specification of R3 and R4, or the formation of symmetric ommatidia, where both cells acquire either the R3 or the R4 fate (Cooper and Bray, 1999; Fanto and Mlodzik, 1999; Strutt et al., 2002; Tomlinson and Struhl, 1999; Zheng et al., 1995). R3/R4 specification is in part mediated by the transcriptional upregulation of the Notch ligand *Delta* and its positive regulator *neuralized* in R3, leading to activation of Notch itself in R4 (Cooper and Bray, 1999; Fanto and Mlodzik, 1999) (del Alamo and Mlodzik, 2006).

Numb is an important negative regulator of Notch signaling during the asymmetric cell divisions that give rise to the cell lineage of the sensory organ precursors and other cell specification events (Uemura et al., 1989) (Rhyu et al., 1994) (Frise et al., 1996) (Buescher et al., 1998) (Carmena et al., 1998). Numb (and alpha-adaptin) act by regulating the endocytosis/recycling of Notch and Sanpodo (Berdnik et al., 2002) (O’Connor-Giles and Skeath, 2003) (Hutterer and Knoblich, 2005) (Couturier et al., 2013) (Johnson et al., 2016). Here, we focus on the role of Numb as part of the molecular mechanisms that regulate the bias of Notch signaling activation in R3/R4 and the establishment of planar cell polarity in the *Drosophila* eye.

## MATERIALS AND METHODS

### Fly stocks and mosaic analysis

The following transgenic and mutant fly stocks were used: *ft^fd^* (Bryant et al., 1988), *E(spl)mδ0.5* (Cooper and Bray, 1999), *numb^15^* (Berdnik et al., 2002), *numb^1^* (BDSC 4096), *numb^2^* (Uemura et al., 1989), *numb4* (Skeath and Doe, 1998), *P(PZ)numb^03235^* (BDSC 11278), *P(EPgy2)numb^EY03840^,FRT40A* (Kyoto 114573), *sev-GAL4* (Richardson et al., 1995), *UAS-Numb*(Frise et al., 1996), *UAS-Numb-GFP* (Song and Lu, 2012), *UAS-CD8-GFP* (BDSC 5136), *dsh^3^,FRT19A* (BDSC 6331), *dsh^1^* (BDSC 5298), *lgl^4^* (Gateff and Schneiderman, 1974), *lgl^4w3^* (Bilder et al., 2000), *tubGAL80,FRT40*A (BDSC 5192), *Diap1-lacZ* (*P(lacW)th^j5C8^*(Ryoo et al., 2002)).

Clones of mutant eye tissue were generated by the Flp/FRT technique (Golic, 1991). Flipase expression was induced under the control of the *eyeless* promoter (Newsome et al., 2000).

### Immunofluorescence and tangential plastic sections

Third instar larval eye discs were dissected and processed for immunofluorescence as described in (Pocas et al., 2015). Briefly, the discs were dissected in 1xPBS, fixed in 1xPBS + 4% Formaldehyde for 20 minutes at room temperature and washed 3 times with PBX (1xPBS + 0.3% Triton x-100). Primary antibodies were incubated in BNT (1xPBS, 1% BSA, 0.1% Tween 20, 250 mM NaCl) overnight at 4°C. Primary antibodies were as follows: anti-β-gal (Cappel), rat anti-ELAV (DSHB), rabbit anti-BarH1 (Higashijima et al., 1992), rabbit anti-Salm (Kuhnlein et al., 1994), mouse anti-γ-tubulin (Sigma GTU-88), rabbit anti-Numb and mouse anti-Fat. Samples were washed 3 times with PBX and incubated with appropriate secondary antibodies (Cy3, Cy5, FITC from Jackson Immuno-Research Laboratories) for 2 hours at room temperature. Samples were mounted in Vectashield (Vector Laboratories) and analyzed on a Zeiss LSM 510 confocal microscope. Tangential sections of adult eyes were performed as described in (Tomlinson and Ready, 1987).

## RESULTS AND DISCUSSION

### Numb is expressed in R3/R4 and upregulated in R3 during PCP establishment

In 3^rd^ instar larval eye imaginal discs, *numb* expression, as assayed by the *P(PZ)numb^03235^* transgenic line, was detected in the R3/R4 pair (and other photoreceptor cells - Fig. 1C), coincident with Salm (Spalt major) expression in these cells, which corresponds to the stage when R3/R4 specification and PCP establishment occur (Domingos et al., 2004a) (Domingos et al., 2004b).

While we detected no obvious difference in the levels of expression of *P(PZ)numb^03235^* between the R3 and R4 cells, using an antibody recognizing Numb, we observed higher levels of membrane associated Numb immunoreactivity in R3 as compared to R4 (arrowheads in Fig. 1D). To confirm these results we used a Numb-GFP fusion protein, under the control of *sev-GAL4*, which revealed an marked upregulation in R3 cells (arrows in Fig.2A), concomitant with *E(spl)mδ0.5* expression in R4, a read out of Notch signaling activation. This upregulation of Numb-GFP in R3 is specific, and was not observed with a control GFP fusion protein, CD8-GFP under *sev-GAL4* control (Fig. 2B). These results exclude the possibility that *sev-Gal4* has stronger expression in R3 than R4, and confirm that the upregulation of Numb occurs via post-transcriptional mechanism. Interestingly, we did not observe an upregulation of Numb-GFP (under *sev-GAL4* control) in null clones of *dishevelled* (*dsh^3^*) (Fig. 2C), indicating that the post-transcriptional upregulation of Numb in R3 is regulated by the Fz-Dsh core PCP pathway.

**Figure 2 –.**
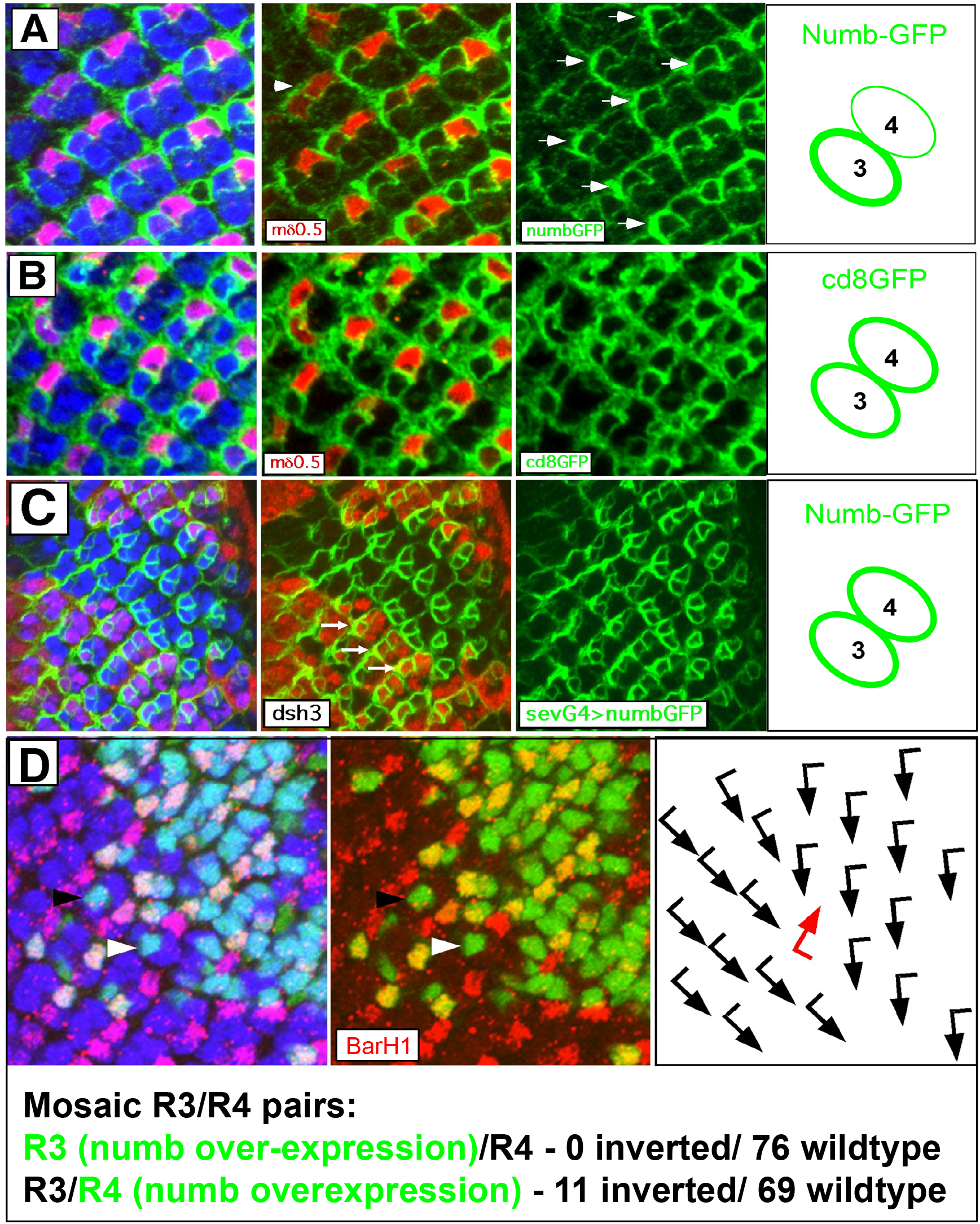
Higher levels of Numb are associated with R3 cell fate. A) Photoreceptors expressing Numb-GFP (green) under the control of sev-Gal4 show higher levels of Numb-GFP in R3 (arrows), transiently, when *mδ0.5-lacZ* (red) is upregulated in R4. Similar levels of numb-GFP in R3/R4 are observed in ommatidia where *mδ0.5-lacZ* (red) is not yet upregulated in R4 (arrowhead) and in more posterior ommatidia (to the right), where R3/R4 fate and ommatidial rotation are resolved. B) Photoreceptors expressing CD8-GFP (green) under the control of sev-Gal4 show identical levels of CD8-GFP in the R3/R4 pair, before, during and after *mδ0.5-lacZ* (red) upregulation in R4. C) Ommatidia homozygous mutant for dsh3, labeled by de absence of DsRed, show identical levels of Numb-GFP in the R3/R4 pair (under the control of sev-Gal4), in contrast to wild-type tissue where Numb-GFP is upregulated in R3 (arrows). D) Overexpression of Numb in R4 leads to chiral inversions. Mosaic overxpression of Numb was done by using larva with the genotype *eyFlp; tubGal80,FRT40A/ubiGFP,FRT40A; sev-Gal4/UAS-Numb*, so that Numb is only overexpressed after Flp induced recombination, in cells containing 2 copies of *ubiGFP,FRT40A*. The black arrowhead indicates an example of an ommatidia of correct chirality, where Numb is overexpressed in R3, but not in R4. The white arrowhead indicates an example of an ommatidia with inverted chirality, where Numb is overexpressed in the presumptive R4, which was transformed into the R3 fate. In 76 ommatidia overexpressing Numb in R3, all present correct chirality. In 69 ommatidia overexpressing Numb in R4, 11 ommatidia had inverted chirality.

### Mosaic overexpression of Numb results in PCP defects

Numb was previously described as an inhibitor of Notch signaling during the differentiation of cell lineages of the sensory organ precursors/SOPs (Frise et al., 1996; Guo et al., 1996), that differentiate into the distinct cell types of the bristle mechanosensory organs of the adult fly. Our observation concerning the upregulation of Numb in R3 is consistent with the role of Numb as negative regulator of Notch signaling, since Notch signaling needs to be repressed in R3, besides being activated in R4 during PCP establishment (Cooper and Bray, 1999; Fanto and Mlodzik, 1999; Tomlinson and Struhl, 1999).

To further explore the role of Numb in R3/R4 specification and PCP establishment, we performed mosaic overexpression experiments with Numb (Fig.2D), using the genotype *eyFlp; tubGal80, FRT40A/ubiGFP, FRT40A; sev-Gal4/UAS-Numb*, so that Numb is overexpressed only after Flp-induced mitotic recombination, in cells containing 2 *ubiGFP, FRT40A* chromosomes but not *tub-GAL80*, an inhibitor of GAL4 activity. Using immunofluorescence assays in the larval eye disc with an antibody against BarH1, we scored the ommatidial rotation phenotypes of mosaic ommatidia, where Numb was overexpressed in R3 but not R4. We detected no rotation defects in ommatidia of such genotype (n=76; Fig. 2). In contrast, when Numb was overexpressed in R4 only, we observed ommatidia (11/80) that inverted the direction of rotation (one example is indicated with an arrowhead in Fig. 2D), indicative of an inverted chirality and hence an inversion in the R3/R4 cell fate decision. In control genotypes, using the same genetic combination but without UAS-Numb (*eyFlp; tubGal80, FRT40A/ubiGFP, FRT40A; sev-Gal4/+*), we did not observe any rotation/chirality defects (data not shown) neither in 62 ommatidia with “green” R3 and “black” R4, nor in 60 ommatidia with “black” R3 and “green” R4, with GAL4 being active in the “green” but not the “black” cells. These results demonstrate that Numb can be instructive and can repress the R4 fate, presumably by inhibiting Notch signaling and by forcing the R4 precursor into an R3 fate.

### Numb levels are regulated by Lgl in the Drosophila larval eye

During the cell divisions of the SOPs, Fz/Dsh signaling regulates cell fate decisions that are determined by the asymmetric localization of Numb (and other cell fate determinants), and the subsequent segregation of these determinants into the daughter cells (Gho and Schweisguth, 1998) (Bellaiche et al., 2001). Our results in the *Drosophila* eye indicate that Fz/Dsh signaling also determines the bias of Numb upregulation in R3 versus R4 (Fig. 2C), despite the fact that the R3/R4 precursors at this stage of PCP establishment are two postmitotic cells, specified by mechanisms that do not involve the asymmetric localization of cell fate determinants during cell division.

We next sought to investigate which other factors could be mediating the upregulation of Numb levels in the R3. We focused on Lgl (Lethal giant larvae), which also controls Numb localization during the asymmetric cell divisions of the SOPs (Langevin et al., 2005). We employed two different *lgl* alleles, *lgl^4^* and *lgl^4w3^*, in clonal mosaic analyses in the *Drosophila* eye, and observed that *sevGAL4>Numb-GFP* levels were generally upregulated in *lgl^4w3^* mutant clones (Fig. 3A). This upregulation of Numb-GFP was not only observed in the R3/R4 pair, but in all cells of the eye imaginal disc expressing Numb-GFP. To exclude possible effects of the *lgl* mutation on the activity of *sevGAL4*, we used an antibody against Numb, which also displayed an upregulation of Numb protein in *lgl^4^* mutant clones (Fig. 3B). Importantly, this upregulation of Numb and Numb-GFP in *lgl* mutant cells was observed across the cell (at all levels within the apical-basal axis of photoreceptors), not resulting from a mislocalization of Numb along the apical-basal axis. To confirm these results, we performed Western blots of eye imaginal discs containing “whole eye” *lgl^4^* mutant clones, which displayed an upregulation of Numb as compared to *y w* control discs (Fig. 3C). Taken together, these results indicate that Numb levels are regulated at the post-transcriptional level downstream of Lgl, by mechanisms that could involve the regulation of Numb mRNA stability or Numb protein degradation, for example. Finally, we observed rotation defects (Fig. 3D) and a reduction of *E(spl)mδ0.5* expression levels in *lgl^4^* mutant ommatidia, particularly in clusters closer to the morphogenetic furrow, when PCP establishment takes place (at the onset of *E(spl)mδ0.5* in R4).

**Figure 3 -.**
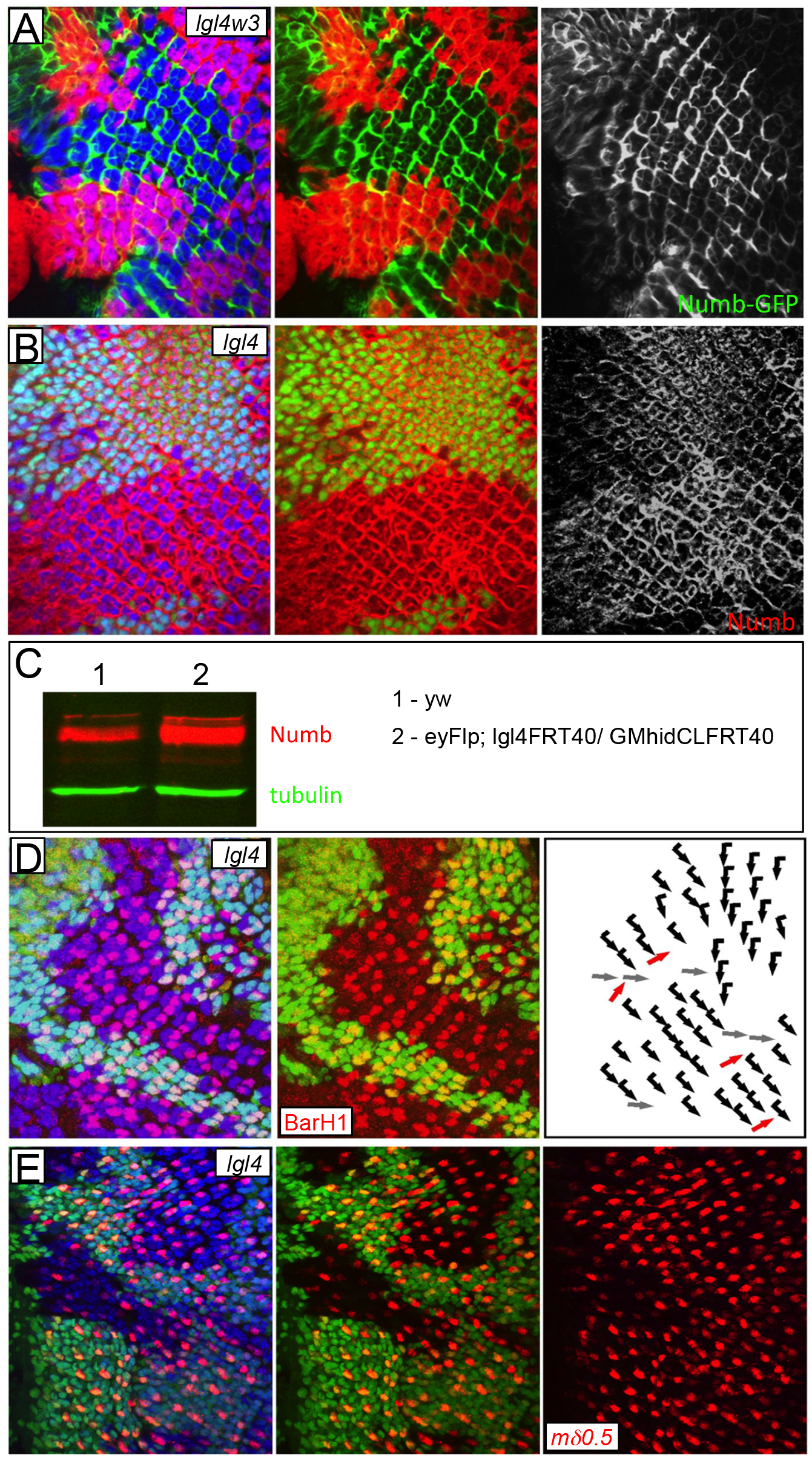
*lgl* mutant clones have higher levels of Numb and PCP defects. A) Numb-GFP (green - under the control of sev-GAL4) is upregulated in *lgl^4w3^* mutant clones, labeled by the absence of DsRed. Elav is in blue. B) Numb (red) is upregulated in *lgl^4^* mutant clones, labeled by the absence of DsRed. Elav is in blue. C) Western blot showing upregulation of Numb in *lgl^4^* “whole eye” mutant clones, dissected from 3^rd^ instar larva eye discs. Tubulin serves as loading control. D) Chiral inversions and rotation defects are observed in *lgl^4^* mutant clones (absence of ubi-GFP). BarH1 is in red and Elav in blue. E) Expression of *m*□*0.5-lacZ* (red) is diminished in *lgl^4^* mutant clones (absence of ubi-GFP), at the onset of R3/R4 specification. Elav is in blue.

### PCP defects in Numb mutants

We next investigated possible PCP defects in several *numb* loss-of-function mutations (Table 1), using immunofluorescence stainings for BarH1 – reflecting ommatidal orientation and hence R3/R4 fate decision - in the larval eye imaginal discs and/or sections of adult eyes. We observed no PCP defects in clones of the *numb^1^* and *numb^2^* alleles (Table 1). Clones of *numb^4^* and *P(EPgy2)numb^EY03840^* showed a low number (around 3%) of mutant ommatidia with PCP defects. Surprisingly, for *numb^15^*, around 20% of mutant ommatidia displayed PCP defects, with chiral inversions being observed both in adults (Fig. 4A) and in larval eye discs (Fig. 4B). *E(spl)mδ0.5* expression was either inverted (arrowheads in Fig. 4C) or presented similar levels of expression in both R3 and R4 (arrows in Fig. 4C) in *numb^15^* mutant clones. Expression of *sevGal4>Numb* caused a partial rescue of the PCP defects, with 6,5% of *numb^15^* mutant ommatidia still presenting rotation inversions (Table 1). Although statistical analyses of the genotypes of the R3/R4 pair in mosaic ommatidia for *numb^15^* (Fig. 5) revealed a requirement for Numb in R3 (Fig. 5A), we also observed sporadic ommatidia with chiral inversions with all photoreceptors being wild-type (example indicated by yellow circle in Fig. 5A). This fact, together with the partial rescue of the PCP defects of *numb^15^* ommatidia by *sevGal4>Numb*, suggested that some other mutation on the *numb^15^* chromosome could be contributing to the strong PCP defects observed.

**Table 1 –.**
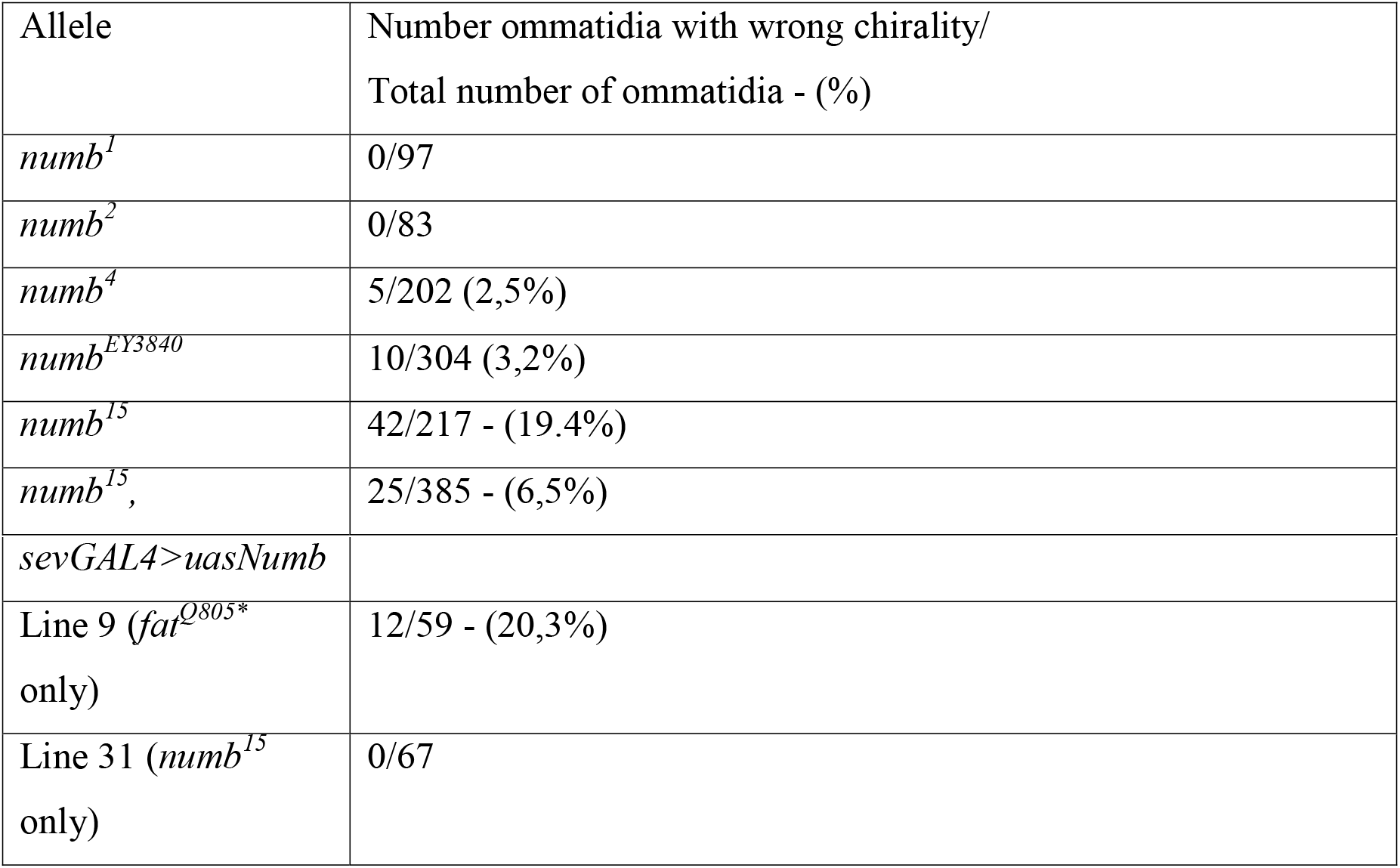
Summary of the chirality inversion phenotypes observed in the indicated alleles and genetic combinations.

**Figure 4 –.**
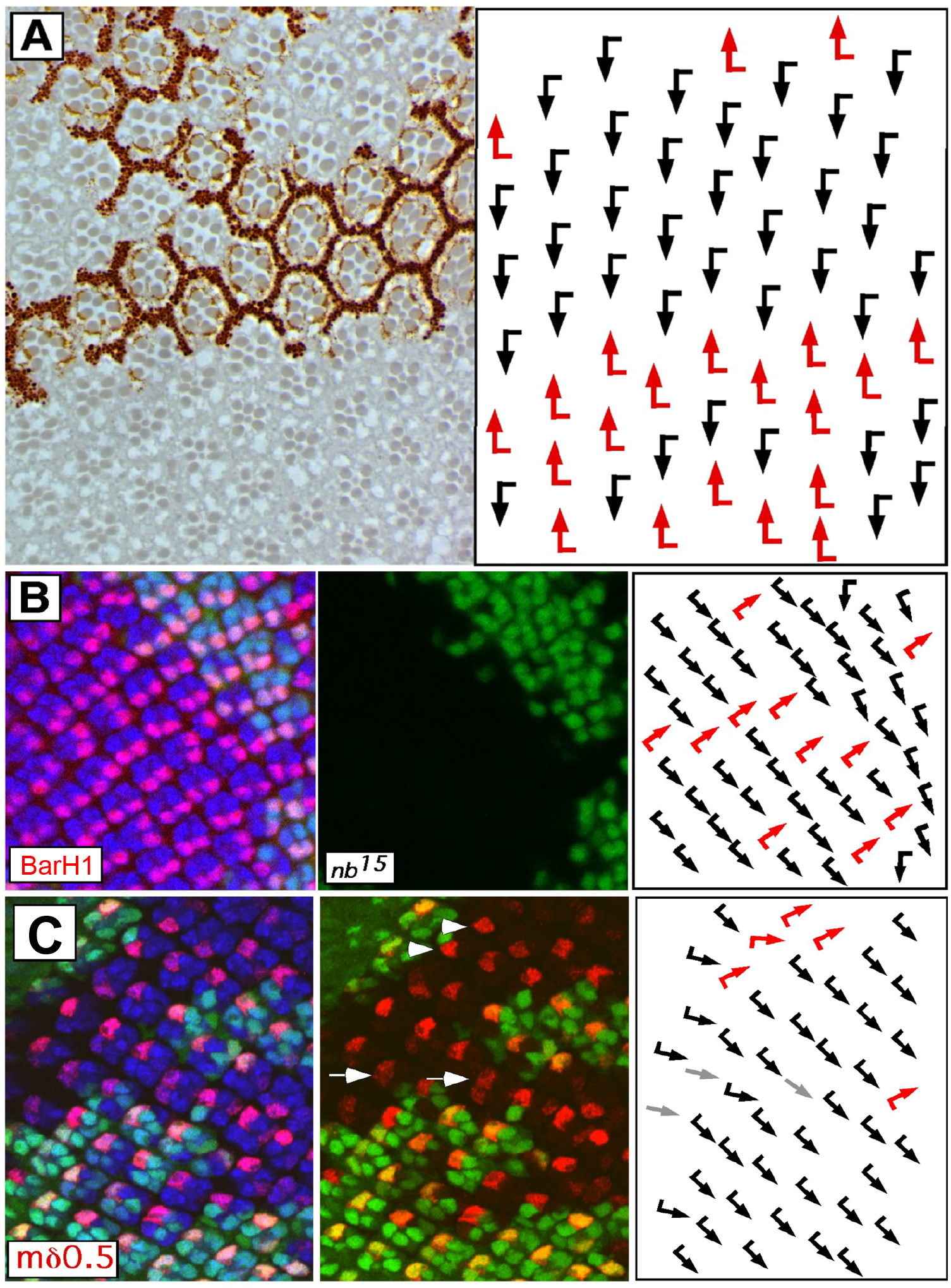
*numb^15^* mutant clones have chiral inversions and rotation defects. A) Tangential sections of adult eyes containing *numb^15^* mutant clones labeled by the absence of the *w*+ marker (detectable as dark dots at the base of each rhabdomere and in pigment cells). Chiral inversions are observed both in the dorsal and ventral halves of the eye. B) Chiral inversions are observed *numb^15^* mutant ommatidia (absence of ubi-GFP) in the larval eye disc. BarH1 is in red and Elav in blue. C) Expression of *mδ0.5-lacZ* (red) is inverted (arrowheads) or persists in both R3 and R4 (arrows) in *numb^15^* mutant ommatidia.

**Figure 5 –.**
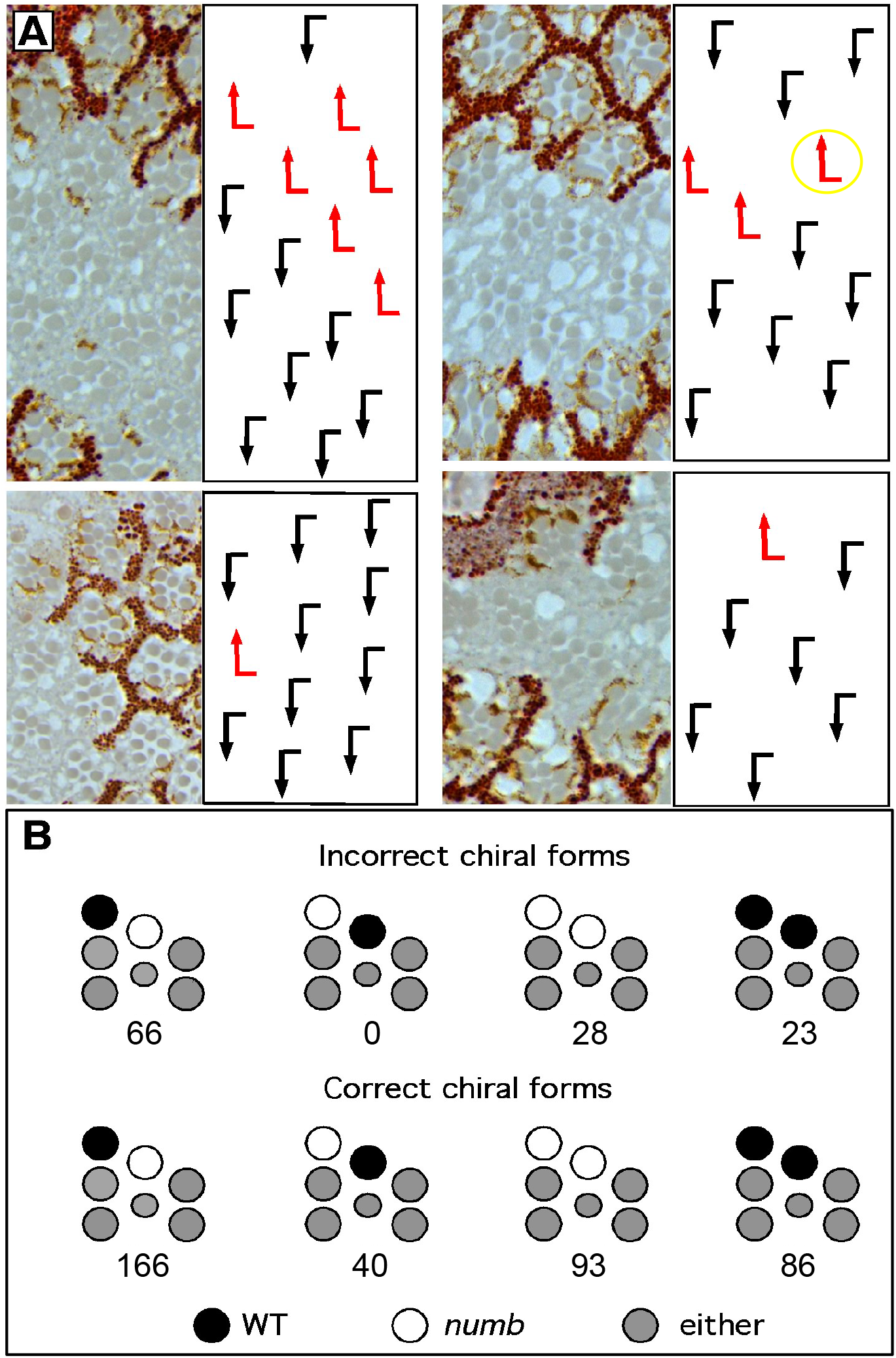
Statistical analysis of *numb^15^* mosaic ommatidia. A) Tangential sections of adult eyes containing *numb^15^* mutant clones presenting ommatidia with mosaic R3/R4 pair. (B) Graphical summary with the statistical analysis of *numb^15^* mosaic ommatidia. *numb^15^* cells are represented as open (white) circles and non-mutant (wild-type-WT) cells as closed (black) circles. As we only scored the genotypes of the R3/R4 pair, other photoreceptors are represented as grey circles (either *numb^15^* or WT). The number of scored ommatidia is indicated below each configuration. For mosaic ommatidia exhibiting chirality inversions, we found 66 ommatidia where R4 has the *numb^15^* genotype and R3 is wild-type. This *numb^15^* mutant R4 cell corresponds to a R3 precursor that made the wrong chiral choice. We have not found any ommatidia with wrong chiral choice where R3 is *numb^15^* and R4 is wildtype. For ommatidia with correct chirality, we found ommatidia with all possible genotypes for the R3/R4 pair.

### The *numb^15^* chromosome contains a non-sense mutation in *fat (ft)*

Based on the nature of the PCP defects that the *numb^15^* chromosome displayed, we hypothesized that it could carry a mutation in the gene encoding the *Fat* protocadherin. *Fat* alleles often are associated with chiral inversions with perfect 90° rotations that follow the wrongly chosen chirality (with no “misrotated” ommatidia with intermediate levels of rotation)(Yang et al., 2002). Also, *fat* resides on the same chromosome arm as *numb* (chromosome 2L), and it is a large gene (with over 15 kb coding the ORF), making *fat* a big target for EMS mutagenesis, that was used to generate *numb^15^* (Berdnik et al., 2002). In fact, we observed that the *numb^15^* chromosome failed to complement a *fat* allele (*fat^fd^*, also known as *fat^8^*), with only 5 escapers reaching adulthood (Table 2), exhibiting a medial cross-vein defect (Fig. 6B) and PCP defects in the eye (Fig. 6C). In addition, we detected a strong reduction of Fat protein levels in *numb^15^* mutant clones (Fig. 6D). We thus sequenced the *fat* locus on the *numb^15^* chromosome and discovered several nucleotide changes leading to amino acid polymorphisms and one premature stop codon, Q805* (Table 3).

**Table 2 –.** *fat^fd^* complements all numb mutations tested except *numb^15^*. The number of adult flies obtained for each cross is presented both for transheterozygotes (non-CyO) and each mutation over the balancer chromosome, CyO.

|  | *fat*<sup>fd</sup>/CyO |
| --- | --- |
| *numb*<sup>15</sup>/CyO | 90 CyO<br>5 non-Cyo (escapers) |
| *numb*<sup>2</sup>/CyO | 45 CyO<br>42 non-CyO |
| *numb*<sup>4</sup>/CyO | 69 CyO<br>28 non-CyO |
| *numb*<sup>1</sup>/CyO | 63 CyO<br>39 non-CyO |
| *numb*<sup>EY3840</sup>/CyO | 45 CyO<br>49 non-CyO |

**Figure 6 –.**
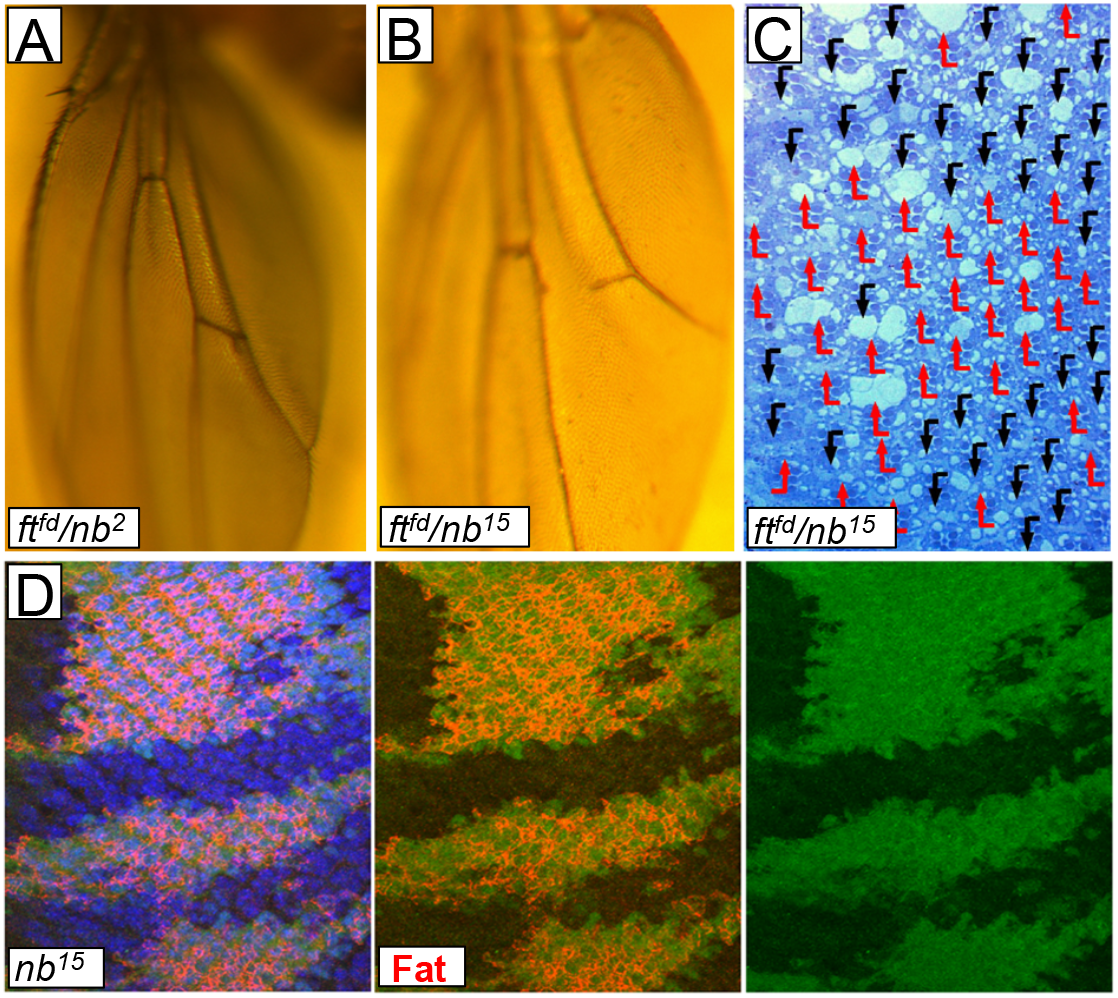
The *numb^15^* chromosome contains a mutation in *fat*. A) Normal adult wing of a trans-heterozygotes of *fat^fd^* and *numb^2^*. B) Medial cross-vein defect in a wing trans-heterozygote for *fat^fd^* and *numb^15^*. C) Adult eye chirality inversion from a transheterozygote of *fat^fd^* and *numb^15^*. D) Larval eye discs containing *numb^15^* mutant clones (absence of ubi-GFP) have reduced levels of Fat (red). Elav is in blue.

**Table 3 –.**
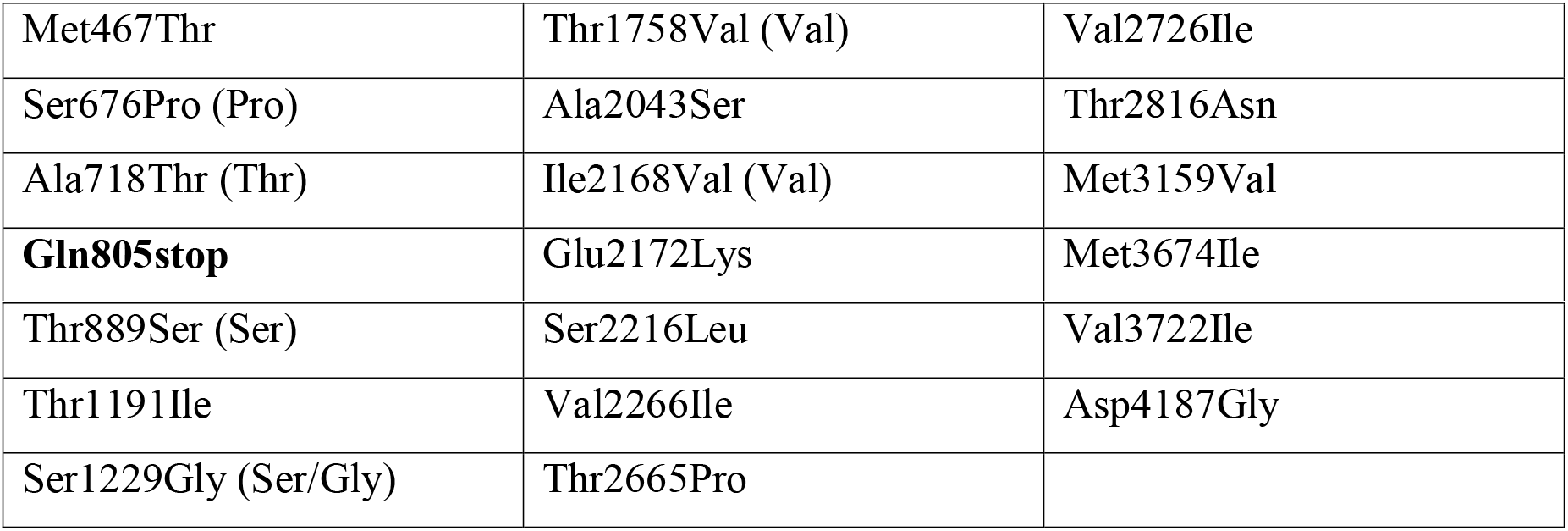
List of mutations found in the *fat* locus in the chromosome containing *numb^15^*. Sequence comparison was made in relation to the sequence available in the public databases (Flybase). In brackets (), is the amino acid found in the sequenced published by (Mahoney et al., 1991).

Subsequent recombination experiments to separate *numb^15^* from *fat^Q805*^*, isolated line 9 that contains *fat^Q805*^* but not *numb^15^* (Fig. 7A-B) and line 31 that contains *numb^15^* but not *fat^Q805*^* (Fig. 7C-D). We detected PCP defects in line 9, but not in line 31 (Table 1), indicating that the PCP defects detected with the original *numb^15^* chromosome were due to *fat^Q805*^*. Premature stop codons, such as *fat^Q805*^*, are usually strong protein null alleles due to the degradation of the mRNA by “Non-Sense mediated RNA decay” (Celik et al., 2015). The PCP defects observed in *fat^Q805*^* are strong and consistent with *fat^Q805*^* being a strong loss-of-function allele. Clones of *fat* mutant cells also behave as supercompetitors, proliferating and outcompeting neighboring non *fat* mutant cells (Tyler and Baker, 2007) (Tyler et al., 2007), and we could observe this supercompetitor behavior of *ft^fd^* mutant cells, that proliferate and completely outcompete *w+* (non mutant) cells in adult eyes (Fig 7E). For *fat^Q805*^* mutant cells, however, the overproliferation phenotype was weaker than in *ft^fd^*, since clones of *w+* cells were still observed in adult eyes (Fig. 7F). Also, we observed *fat^Q805*^* mutant cells undergoing apoptosis, assayed by immunofluorescence staining with an antibody against active caspase 3 (Fig. 7H), which was not observed in *ft^fd^* mutant clones (Fig. 7G). Finally, while *ft^fd^* mutant cells upregulate the expression of Diap1-lacZ (Fig. 7I), as previously shown for another strong *fat* allele (Tyler and Baker, 2007), *fat^Q805*^* mutant cells fail to do so (Fig 7J). These results indicate that the *fat^Q805*^* mutation is of hypomorphic nature regarding overproliferation phenotypes, which is surprising, given the molecular basis of *fat^Q805*^* (a premature stop codon) and the strong PCP defects phenotypes observed.

**Figure 7 –.**
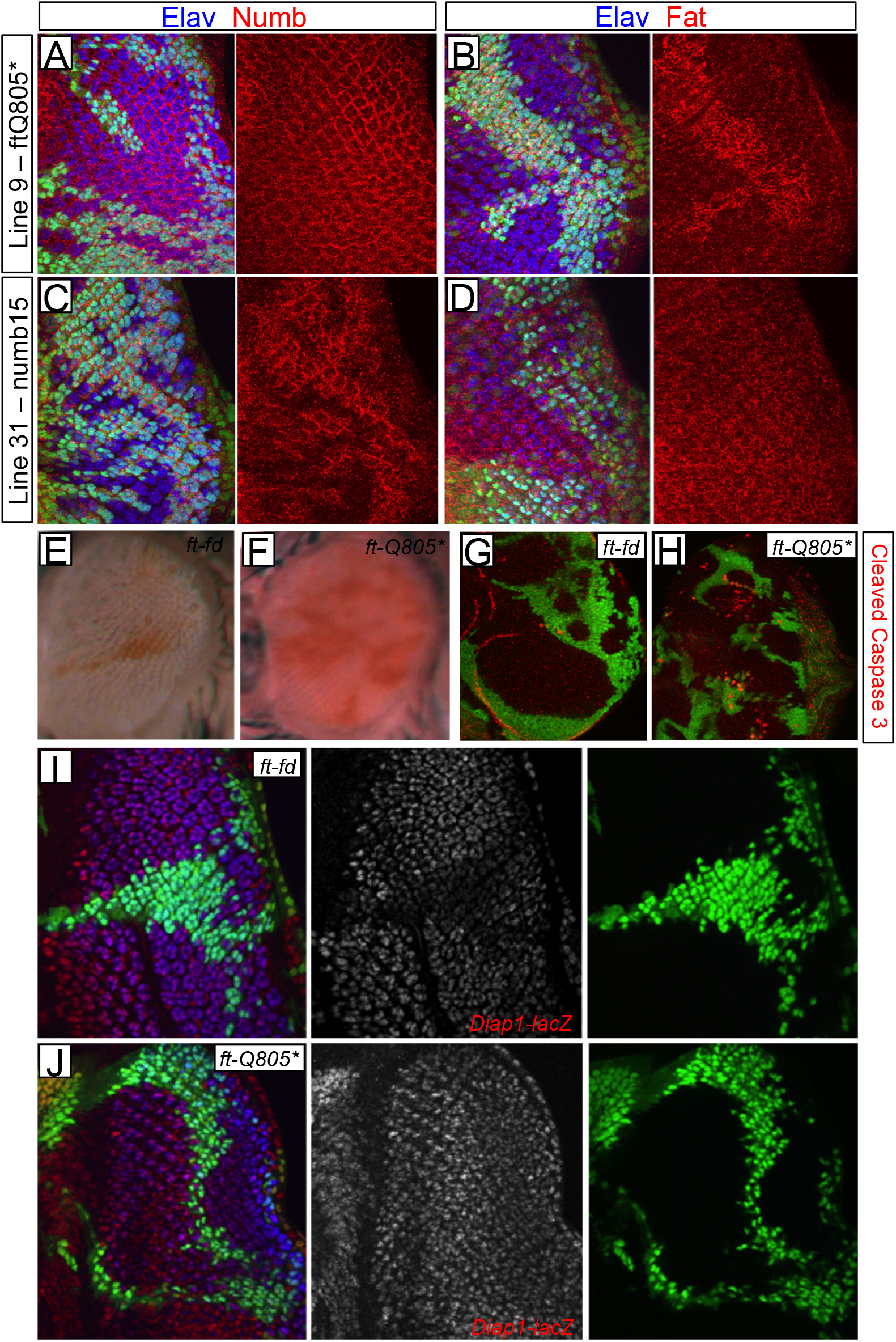
Characterization of *fat^Q805*^* mutation. A) Eye imaginal disc containing clones of “line 9” (absence of ubi-GFP), a chromosome containing *fat^Q805*^* but not *numb^15^* (obtained from recombination of the original *numb^15^* chromosome) presents normal levels of Numb (red). Elav is in blue in all panels. B) Clones of “line 9” have reduced levels of Fat (red). C) Clones of “line 31” (a chromosome containing *numb^15^* but not *fat^Q805*^* have reduced levels of Numb (red). D) Clones of “line 31” have normal levels of Fat. E) Adult eye containing clones of *fat^fd^*, labeled by the absence of w+. F) Adult eye containing clones of *fat^Q805*^*, labeled by the absence of w+. G) Eye imaginal disc containing clones of ft-fd (absence of ubi-GFP) have low levels of active caspase 3 (red). H) Eye imaginal disc containing clones of *fat^Q805*^* (absence of ubi-GFP) present cells undergoing apoptosis (active caspase 3 - red). I) Eye imaginal disc containing clones of *fat^fd^* (absence of ubi-GFP) have upregulated levels of *Diap1-lacZ* (red and monochrome). J) Eye imaginal disc containing clones of *fat^Q805*^* (absence of ubi-GFP) have normal levels of *Diap1-lacZ* (red and monochrome).

### *numb* acts redundantly in the PCP-mediated R3/R4 specification

A surprising observation was the fact that *sevGAL4>Numb* caused a significant rescue of the original *numb^15^* chromosome (Table 1), even if partial, of the PCP defects observed in that *numb^15^* chromosome, given that these PCP defects were caused by *fat^Q805*^*. We cannot offer a definitive explanation for this finding. However, our data strongly suggest that Numb is nevertheless involved in the R3/R4 cell fate decisions and PCP signaling associated process. Both the gain-of-function (*sevGAL4>Numb*) and associated mosaic overexpression of Numb in the R3/R4 pair modified R3/R4 cell fates, and *numb^2^* mutant chromosomes dominantly suppressed the PCP defects in the *dsh^1^* mutation (Table 4). Moreover, the upregulation of Numb protein levels in R3 also depended on Fz/Dsh PCP signaling, as it was lost in the *dsh* null allele, *dsh^3^*. Taken together, these data suggest that Numb contributes to the PCP-associated R3/R4 specification outcome, albeit through a redundant mechanism. Modifying Numb levels, by overexpression, or by using a *numb* mutant background in sensitized PCP genotypes, as observed for *fat^Q805*^* or *dsh^1^* backgrounds revealed a limitation in the correct fate decisions within the R3/R4 cell pair, thus revealing a role for Numb in this process.

**Table 4 –.** Summary table of the chirality and rotation defects found in males *dsh^1^*/Y and *dsh^1^*/Y; *numb^2^*/+. *dsh^1^* is an hypomorphic mutation presenting chiral and rotation defects and symmetric ommatidia with R3/R3 or R4/R4 configurations.

|  | Normal Chirality | Inverted Chirality or misrotated | R3/R3 or R4/R4 |
| --- | --- | --- | --- |
| *dsh<sup>1</sup>/Y* | 211 (66,3%) | 36 (11,6%) | 71 (22,3%) |
| *dsh<sup>1</sup>/Y; numb<sup>2</sup>/+* | 444 (87,1%) | 46 (9,0%) | 20 (3,9%) |

## Acknowledgements

We thank Seth Blair, Bingwei Lu, Sarah Bray, the Bloomington Drosophila Stock Center at the University of Indiana and the Developmental Studies Hybridoma Bank at the University of Iowa for fly strains and antibodies. We are grateful to Jurgen Knoblich for the *numb^15^* allele and discussions about our results. We thank François Schweisguth for helpful discussions. We thank Keon Combie for assistance in sequencing the *fat* mutant allele. H.S. was an Investigator with the Howard Hughes Medical Institute. This work was supported by the Strang Foundation and NIH grants from the National Eye Institute (R01 EY14025) to BM and (RO1 EY13256) to MM. Additional funding was granted to PMD (FCT LISBOA-01-0145-FEDER-007660, PTDC/NEU-NMC/2459/2014 and IF/00697/2014) and to BM (ANR 13-ISV4-0003-01).

